# A strain of an emerging Indian pathotype of *Xanthomonas oryzae* pv. oryzae defeats the rice bacterial blight resistance gene *xa13* without inducing a clade III *SWEET* gene and is nearly identical to a recent Thai isolate

**DOI:** 10.1101/384289

**Authors:** Sara C. D. Carpenter, Prashant Mishra, Chandrika Ghoshal, Prasanta Dash, Li Wang, Samriti Midha, Gouri S. Laha, Jagjeet S Lore, Wichai Kositratana, Nagendra K. Singh, Kuldeep Singh, Prabhu B. Patil, Ricardo Oliva, Sujin Patarapuwadol, Adam J Bogdanove, Rhitu Rai

**Affiliations:** Plant Pathology and Plant-Microbe Biology section, School of Integrative Plant Science, Cornell University, Ithaca, New York, USA; Plant Pathogen Interaction, ICAR-National Research Centre on Plant Biotechnology, Pusa, New Delhi, India; CSIR-Institute of Microbial Technology, Chandigarh, India; ICAR-Indian Institute of Rice Research, Rajendranagar, Hyderabad, Telangana, India; Department of Plant Pathology, Punjab Agricultural University, Ludhiana, Punjab, India; Center for Agricultural Biotechnology, Kamphaeng Saen, Kasetsart University, Kamphaeng Saen Campus, Nakhon Pathom, 73140, Thailand; ICAR-National Bureau of Plant Genetic Resources, New Delhi, India; International Rice Research Institute, Los Banos, Philippines

**Keywords:** bacterial blight of rice, SMRT sequencing, transcription activator-like effectors (TALEs), susceptibility genes, *SWEET* genes

## Abstract

The rice bacterial blight pathogen *Xanthomonas oryzae* pv. oryzae (*Xoo*) injects transcription activator-like effectors (TALEs) that bind and activate host ‘susceptibility’ (*S*) genes important for disease. Clade III *SWEET* genes are major S genes for bacterial blight. The resistance genes *xa5*, which reduces TALE activity generally, and *xa13*, a *SWEET11* allele not recognized by the cognate TALE, have been effectively deployed. However, strains that defeat both resistance genes individually were recently reported in India and Thailand. To gain insight into the mechanism(s), we completely sequenced the genome of one such strain from each country and examined the encoded TALEs. Strikingly, the two strains are clones, sharing nearly identical TALE repertoires, including a TALE known to activate *SWEET11* strongly enough to be effective even when diminished by *xa5*. We next investigated *SWEET* gene induction by the Indian strain. The Indian strain induced no clade III *SWEET* in plants harbouring *xa13*, indicating a pathogen adaptation that relieves dependence on these genes for susceptibility. The findings open a door to mechanistic understanding of the role *SWEET* genes play in susceptibility and illustrate the importance of complete genome sequence-based monitoring of *Xoo* populations in developing varieties with effective disease resistance.

## Introduction

*Xanthomonas oryzae* pv. oryzae (*Xoo*), causes bacterial blight of rice, a yield-reducing disease widespread in Asia and Africa ^1^. *Xoo* relies on type III secreted, transcription activator-like effectors (TALEs) that directly activate specific host genes, called ‘susceptibility’ (*S*) genes, which contribute to disease development ^2^. A TALE finds its DNA target by virtue of a central repeat region (CRR) in the protein composed of nearly identical, direct repeats of 33-35 amino acid residues. Residues at the 12^th^ and 13^th^ positions in each repeat, together the “repeat-variable diresidue” (RVD), correspond to a single nucleotide in the effector binding element (EBE) in the DNA in a contiguous, code-like fashion such that the number and composition of RVDs predict the sequence of the EBE ^3,4^. The first residue of each RVD plays a stabilizing role and the second is the base-specifying residue. Characterized *Xoo* strains harbour 9 to nearly 20 different TALE-encoding (*tal*) genes, of which only one or two may encode a major virulence factor ^2,5,6^. All strains examined to date activate one of three members of clade III of the *SWEET* sucrose transporter gene family in rice (*SWEET11*, *SWEET12*, and *SWEET14*). These genes are major *S* genes, targeted by diverse TALEs from different strains ^2^. In an experimental context, each of the other two members of *SWEET* clade III (*SWEET12* and *SWEET15*), and no other *SWEET* genes tested, also functioned as a major *S* gene ^7,8^. *SWEET* activation apparently leads to sucrose export into the xylem vessels, facilitating *Xoo* proliferation and symptom development by an as yet uncharacterized mechanism.

Host resistance is the most effective means of controlling rice bacterial blight. To date, 42 bacterial blight resistance genes, called *Xa* genes, have been identified from cultivated and wild rice species ^2,9,10^. The functions of most of the dozen or so that have been cloned and characterized relate to TALEs, and several are recessive. All but one of these recessive genes are alleles of a *SWEET* gene with a mutation at the EBE that prevents binding and activation by the cognate TALE, conferring resistance through reduced susceptibility. For example, *xa13* is a variant of *SWEET11* that lacks the PthXo1 EBE in its promoter and thereby confers resistance to strains that depend on PthXo1 ^11^. A strain can overcome *xa13* if it expresses a TALE (such as PthXo2, PthXo3, AvrXa7, or TalC) that activates an alternate clade III *SWEET* gene ^12^. The recessive bacterial blight resistance gene that is not a *SWEET* allele, *xa5*, acts more broadly. It is an allele of the general transcription factor subunit gene *TFIIAγ5*. The protein encoded by the dominant allele is an apparent contact point between TALEs and the transcriptional machinery. The product of *xa5* harbours a single amino acid substitution that interferes with its interaction with TALEs and thereby reduces activation of their targets ^13,14^. Interestingly, strains carrying PthXo1 are compatible with *xa5*. This compatibility is postulated to be due to the unusually strong activation of *SWEET11* by PthXo1, which even diminished in the *xa5* background is apparently high enough to render the plant susceptible ^13^. The *xa5* and *xa13* genes have been widely deployed, both singly and in combination ^15–18^. Their effectiveness, however, has varied in different rice growing countries. In India, which is the second largest producer of rice behind China and has a highly diverse *Xoo* population ^19^, *xa13* has historically been effective, whereas xa5-compatible *Xoo* isolates can be found throughout the country (Figure 1a) ^20,21^. In contrast, in Thailand, another major rice producer, *xa13*-breaking stains are common while *xa5* has largely remained effective (Figure 1b) ^22^. Recently, strains compatible with either *R* gene have been reported in each country ^20–23^. To gain insight into the mechanism(s) by which such strains overcome *xa5* and *xa13*, we completely sequenced and compared the genomes and encoded TALE repertoires of one such strain from each country, IX-280 from India ^21^ and SK2-3 from Thailand ^22^, to each other and to those of other sequenced *Xoo* strains. Further, we examined the ability of the Indian strain to activate *SWEET* gene expression in rice genotypes harbouring *xa5*, *xa13*, or both genes.

**Figure 1:**
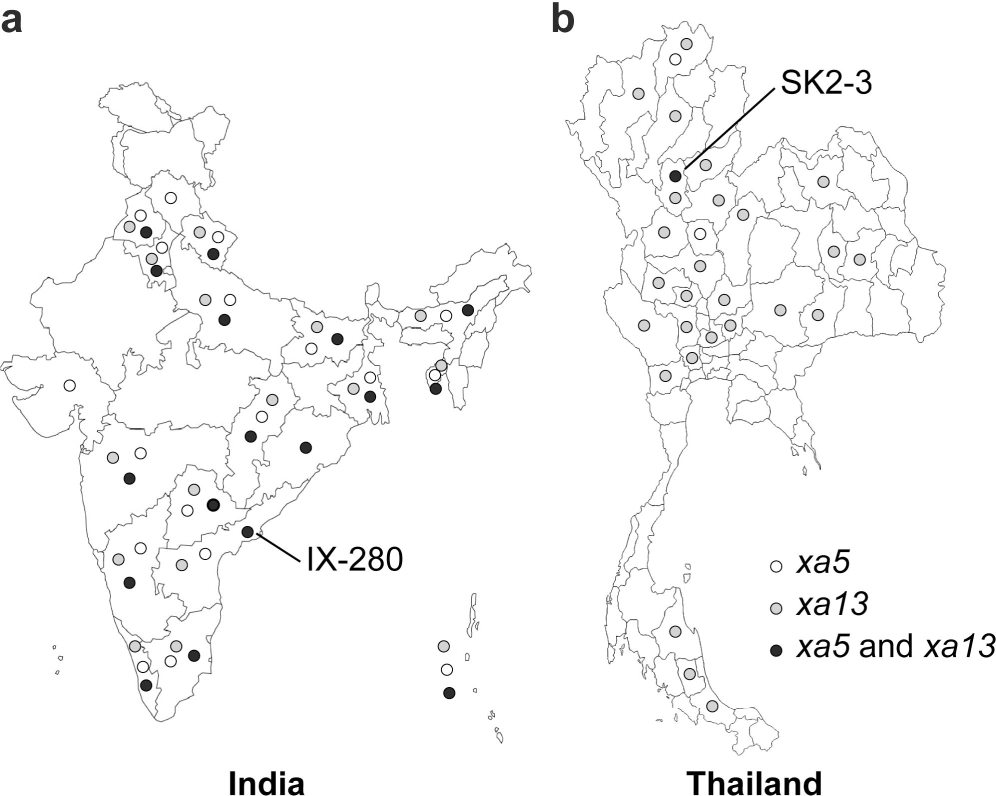
Distribution of *Xoo* strains compatible with rice varieties IRBB5 (*xa5*, white), IRBB13 (*xa13*, grey), and IRBB53 (*xa5*+*xa13*, black) in (a) India and (b) Thailand.

## 1. Materials and Methods

### 1.1. Genomic DNA extraction and sequencing

DNA for complete-genome sequencing was isolated using the protocol described by Booher et al. ^24^ with the following two modifications: after overnight culture and centrifugation, extracellular polysaccharide was removed by washing the bacterial pellet 7-8 times with NE buffer (0.15 M NaCl, 50 mM EDTA), and after cell lysis, DNA was extracted four times with phenol/chloroform and once with chloroform/isoamyl alcohol. For each strain, 4-7 µg of genomic DNA was used to prepare a 20 kb library and each library was sequenced by SMRT technology to >150X genome coverage using P6-C4 chemistry (Pacific Biosciences, Menlo Park, CA USA), as described ^24^.

### 1.2. Genome sequence assembly

*De novo* assembly of the sequence reads was performed using HGAP v.2.0 (HGAP2) and HGAP v. 3.0 (HGAP3) ^25^ as described ^24^. Since TALE encoding (*tal)* genes are often clustered and their repetitive sequences can lead to misassembly even using long-read technology, *tal* gene containing regions were separately assembled using the PBX toolkit, a pipeline that uses long, *tal* gene sequence-containing seed reads to assemble *tal* clusters with more accuracy ^24^. Length cutoff settings used for these seed reads were 16 kb (pbx16000), 12 kb (pbx12000), or 10kb (pbx10000). After HGAP and PBX assemblies were completed, the HGAP assemblies with the fewest unitigs and the majority of the *tal* gene sequences found by PBX were chosen for manual closure and finishing.

### 1.3. Genome finishing, assembly verification, and annotation

To finish the genomes, the circular assemblies were polished twice more with Quiver and then checked for structural variants and misassemblies using PBHoney ^26^. The *tal* gene repertoires were verified by consensus with the local *tal* assemblies made with PBX and by Southern blots of genomic DNA digested with either *Bam*HI or *Sph*I, or with *Bam*HI and *Eco*RI, and probed with the *tal* gene specific probe pZWavrXa7 ^6^. To confirm the absence of plasmids smaller than 20 kb that could have been excluded during library preparation, total DNA was prepared and examined by agarose gel electrophoresis as described, using *Xanthomonas campestris* pv. vesicatoria 85-10, which has four plasmids, as a positive control ^24^. After finishing and assembly verification, genomes were annotated using the NCBI Prokaryotic Genome Annotation Pipeline ^27^, and *tal* gene annotations were manually corrected.

### 1.4. Genomic comparisons

Complete genomes of representative Asian *Xoo* strains were compared using progressiveMauve ^28^ in the MegAlign Pro module of the DNAStar Suite (Lasergene 13.0.0.357) with default settings. For multi-genome phylogenies, all available *Xoo* genomic sequences were aligned using Mauve v2.3.1 ^29^, and core alignment was used to infer phylogeny using PhyML v3.1 ^30^. The core alignment and maximum likelihood tree was further subjected to ClonalFrameML ^31^ analysis with 100 bootstrap replicates to refine the phylogeny considering the impact of recombination. The ClonalFrameML tree was visualized using iTOL v3 ^32^.

### 1.5. TALE analysis and target prediction

All *tal* gene sequences were extracted using the PBX exporter ^24^ or AnnoTALE ^33^. Orthology of IX-280 and SK2-3 TALEs to previously sequenced TALEs was determined using FuncTAL ^34^ and AnnoTALE ^33^. RVD or amino acid sequence was used as input for FuncTAL, and DNA sequence for AnnoTALE. AnnoTALE class builder files used to assign TALEs to families were downloaded on July 1, 2017. The results from the two tools were consistent. Targets of IX-280 and SK2-3 TALEs of interest were predicted using the TALE-NT 2.0 Target Finder tool ^35^ and TalGetter ^36^. Predictions were made for both forward and reverse strands of promoter sequences, defined as the 1000 bp upstream of a transcriptional start site to the translational start site for TALgetter, and 1000 bp upstream of the translational start site for TALE-NT 2.0, and using MSU Rice Genome Annotation Project Release 7 (http://rice.plantbiology.msu.edu/). Default settings were used for Target Finder (upstream base of binding site = T, score cutoff = 3.0, Doyle et al. scoring matrix) and for TALgetter (standard model, p= 0.000001, upstream/downstream offset = 0).

### 1.6. Bacterial and plant growth conditions and disease and gene expression assays

Plants were grown in a growth chamber maintained at 28 °C and 85% relative humidity with a photoperiod of 12 h. The bacterium was cultured at 28° C on modified Wakimoto agar medium. For the disease assay, bacterial cells were resuspended in sterile water at an OD_600_ of 0.2 and clip-inoculated ^37^ to fully expanded leaves of 40-45 day-old plants. Lesions were photographed 14 days later. For gene expression assays, bacterial cells were resuspended in sterile water at an OD_600_ of 0.5 and infiltrated into leaves of three-week-old plants using a needleless syringe. Water was used for mock inoculation as a control. The inoculated portions of leaves were harvested 24 h later, and total RNA was extracted using the PureLink™ RNA Mini kit (Invitrogen, Carlsbad, California, USA) following the manufacturer’s instructions. RNA was further treated with DNase (Invitrogen) to remove genomic DNA contamination. Quality and quantity of RNA were analysed by 1.0% agarose gel electrophoresis and spectrophotometry using a Nanodrop (Thermo Scientific, Waltham, Massachusetts, USA). cDNA was generated from 1 µg purified RNA using the Superscript™ Vilo™ cDNA synthesis kit (Invitrogen) with random primers. Quantitative real time PCR (qPCR) was performed on a Light cycler® 480 Instrument II (Roche Molecular Diagnostics, Santa Barbara, California, USA). About 250 ng of cDNA was used for each qPCR reaction with gene specific primers (Supplementary Table S1). Each gene was tested with three biological replicates, with three technical replicates each. The average threshold cycle (Ct) was used to determine the fold change of gene expression. The expression of each gene was normalized to the expression of the 18S rRNA gene. The 2^ΔΔCt^ method was used for relative quantification ^38^.

## 2. Results

### 2.1. Assembly of the complete IX-280 and SK2-3 genomes

Single Molecule Real-Time (SMRT) DNA sequence data for IX-280 assembled using either HGAP2 or HGAP3 (see Methods) resulted into two contigs, corresponding to a chromosome and a 43 kb plasmid. We named the plasmid pXOO43. The HGAP2 assembly, though it yielded an intact, self-complementary chromosomal contig, collapsed one cluster of four *tal* genes into three, indicated by a coverage spike in that cluster. A comparison of the ends of the misassembled cluster to pbx12000 and pbx16000 assemblies generated using the PBX toolkit ^24^ showed overlap with several that included an intact cluster of four *tal* genes. We chose a pbx16000 contig assembled using settings of 3000 kb read overlap and 97% read identity to replace the misassembled cluster in the HGAP2 assembly. We also verified the presence of the cluster of four *tal* genes in the raw sequence of IX-280. To further confirm our final assembly, we obtained additional long reads from a separate DNA preparation of the same isolate and reassembled with HGAP3 using all available reads; the resulting HGAP3 assembly was consistent with the manually corrected HGAP2 assembly.

HGAP2 and HGAP3 assemblies of SK2-3 yielded a single chromosomal contig, but each terminated at a partial cluster of four *tal* genes. The intact cluster was present in pbx10000 assemblies. We selected a contig assembled using settings of 3000 kb read overlap and 97% read identity to replace the broken cluster in the HGAP2 assembly and manually closed the genome.

The quality-control tool PBHoney ^26^ indicated no major inversions, deletions, or duplications in the assemblies. The proportion of mapped reads to post-filtered reads was 94.9% for IX-280 and 92% for SK2-3. Coverage graphs for the final assemblies showed no unusual peaks or dips that might indicate collapsed or expanded genomic repeats. PBX results were consistent with *tal* gene sequences extracted from the genomes, as were Southern blots hybridized with a *tal* gene-specific probe (Supplementary Figure S1). Separate DNA extraction and gel electrophoresis for both strains confirmed the absence of any small plasmids that might have been missed by SMRT sequencing (not shown).

The IX-280 genome assembly has been deposited in GenBank under accessions CP019226 (chromosome) and CP019227 (plasmid pXOO43) and the SK2-3 assembly under accession CP019515. Raw data and associated metadata are available from the Sequence Read Archive (SRA) under accessions SRR5989134 (IX-280) and SRR5990719, SRR5990720, and SRR5990721 (SK2-3).

### 2.2. Comparison of the IX-280 and SK2-3 genomes

The IX-280 plasmid pXOO43 has not been found in other *Xanthomonas* genomes, but some regions have a high degree of nucleotide identity with regions of pXAC64 from *Xanthomonas citri* ssp. citri ^39^. There are no predicted type III effector genes on the plasmid, but it harbours a cluster of genes annotated as type VI secretion genes. Associated with this cluster is an apparent operon containing *pemK*, encoding a toxin in a toxin/antitoxin system ^40^, and a gene encoding a protein of the XF1863 family, hypothesized to function as its antitoxin ^41^. None of the pXOO43 content is found in the SK2-3 genome.

The IX-280 and SK2-3 chromosomes are entirely syntenous (Figure 2a and Supplementary Figure S2), including the *tal* genes, which show no duplications, deletions, or rearrangements in one genome relative to the other (Figure 2b). To determine how the genome structure of IX-280 and SK2-3 compares with that of other *Xoo* strains, we aligned the genomes with those of completely sequenced strains representing different lineages: PXO99A, PXO86, PXO71 (isolated in the Philippines), MAFF311018 (isolated in Japan), and AXO1947 (isolated in Africa) ^19,42^. The alignment shows no relationship between geographic area of isolation and genome arrangement (Figure 2a). Like IX-280 and SK2-3, the genome structures of PXO71 (Philippines) and MAFF311018 (Japan) are similar to one another, despite the strains being from different countries. In contrast, PXO86, PX071, and PXO99A, all from the Philippines, have undergone genomic rearrangements relative to one another. The genome structure of the African strain, AXO1947, is distinct from those of the other *Xoo* strains, showing some of the genomic variability encompassed by the species. Though there are areas of similarity, the genomic arrangement of IX-280 and SK2-3 is not shared by any of the other strains.

**Figure 2:**
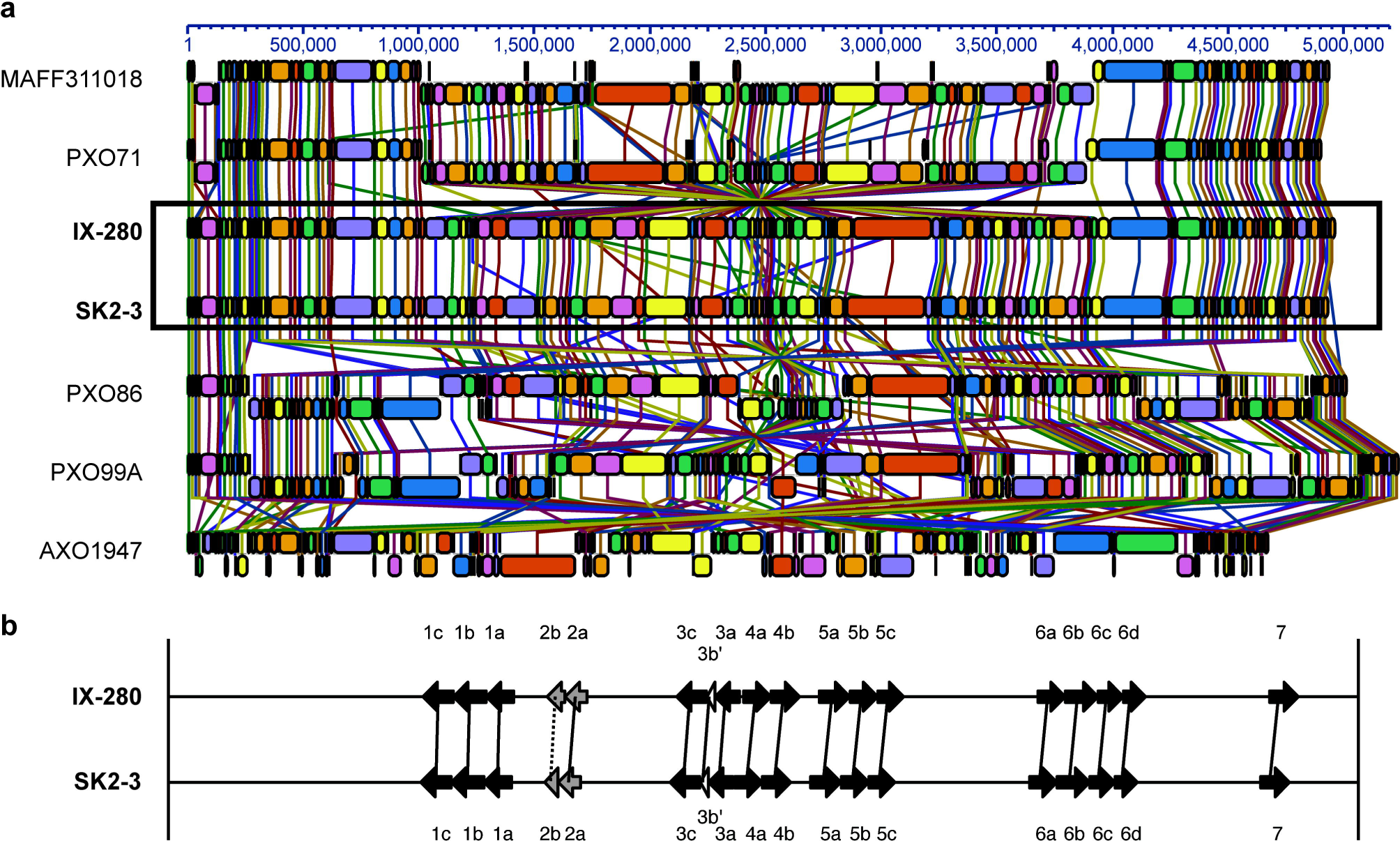
Synteny between IX-280 and SK2-3 genomes and comparison of their *tal* genes. (a) Progressive Mauve alignment of the chromosomes of IX-280 and SK2-3 and other representative *Xoo* strains. (b) Map of the *tal* genes in IX-280 and SK2-3. Black arrows represent full-length *tal* genes, grey arrows truncTALE genes, and white arrows *tal* pseudogenes. Solid lines connect *tal* genes with >99% nucleotide identity and identical RVD sequence, and dotted lines connect less similar but clearly orthologous genes.

### 2.3. IX-280 and SK2-3 belong to a highly clonal lineage

The striking genomic similarity of IX-280 and SK2-3 despite their geographic separation led us to explore their relatedness with other *Xoo* strains more broadly. Using all available *Xoo* complete genomes and draft (short-read derived) genome sequences of 100 Indian *Xoo* strains previously subjected to phylogenetic analysis ^19^, we generated a phylogenetic tree using regions not affected by recombination. Both IX-280 and SK2-3 map to the youngest and a highly clonal lineage, L-I (Figure 3) ^19^. Of the strains examined, SK2-3 is the only non-Indian strain in this lineage.

**Figure 3:**
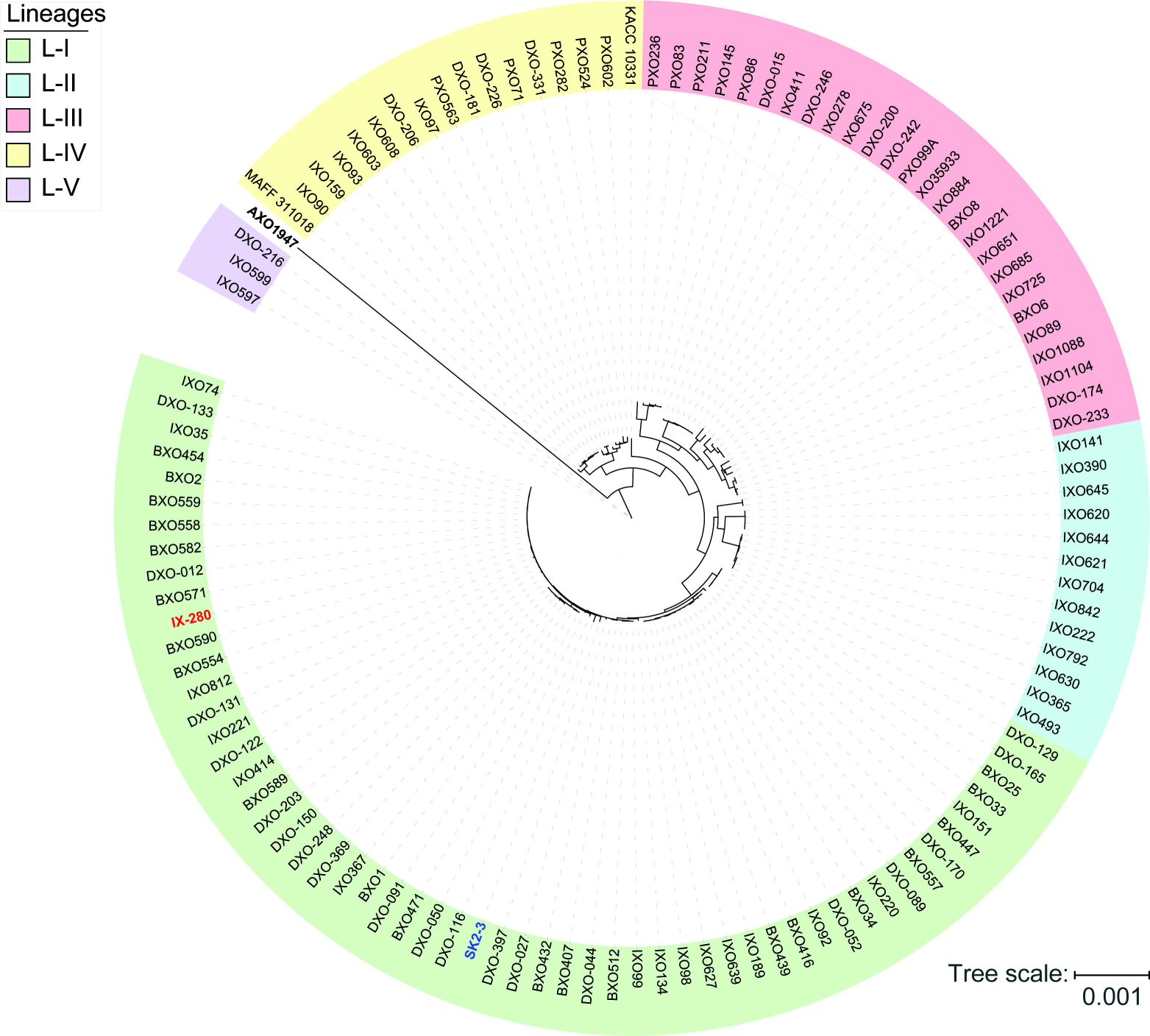
Positions of IX-280 and SK2-3 on a clonal lineage tree derived from genomic sequences of 100 Indian *Xoo* strains and other *Xoo* strains from Asia and Africa. Lineages are block shaded in different colours. IX-280 (red font) and SK2-3 (blue font) are in lineage L-I.

### 2.4. The TALE repertoires suggest possible mechanisms of *xa5* and *xa13* defeat

The TALE repertoires of IX-280 and SK2-3 each consist of 15 TALEs and two truncTALEs, which are TALE variants with shortened N- and C-termini that can function as suppressors of resistance mediated by certain non-executor *R* genes ^43,44^; each strain also harbours a *tal* pseudogene (Figure 4). The RVD sequence of each IX-280 TALE and truncTALE is identical to that of its counterpart in SK2-3, except for the truncTALE Tal2b, of which repeats 10-15 are missing in SK2-3. Since truncTALEs do not bind DNA and a specific RVD sequence is not critical to their function ^43^, this difference in Tal2b between the two strains is likely functionally irrelevant.

**Figure 4:**
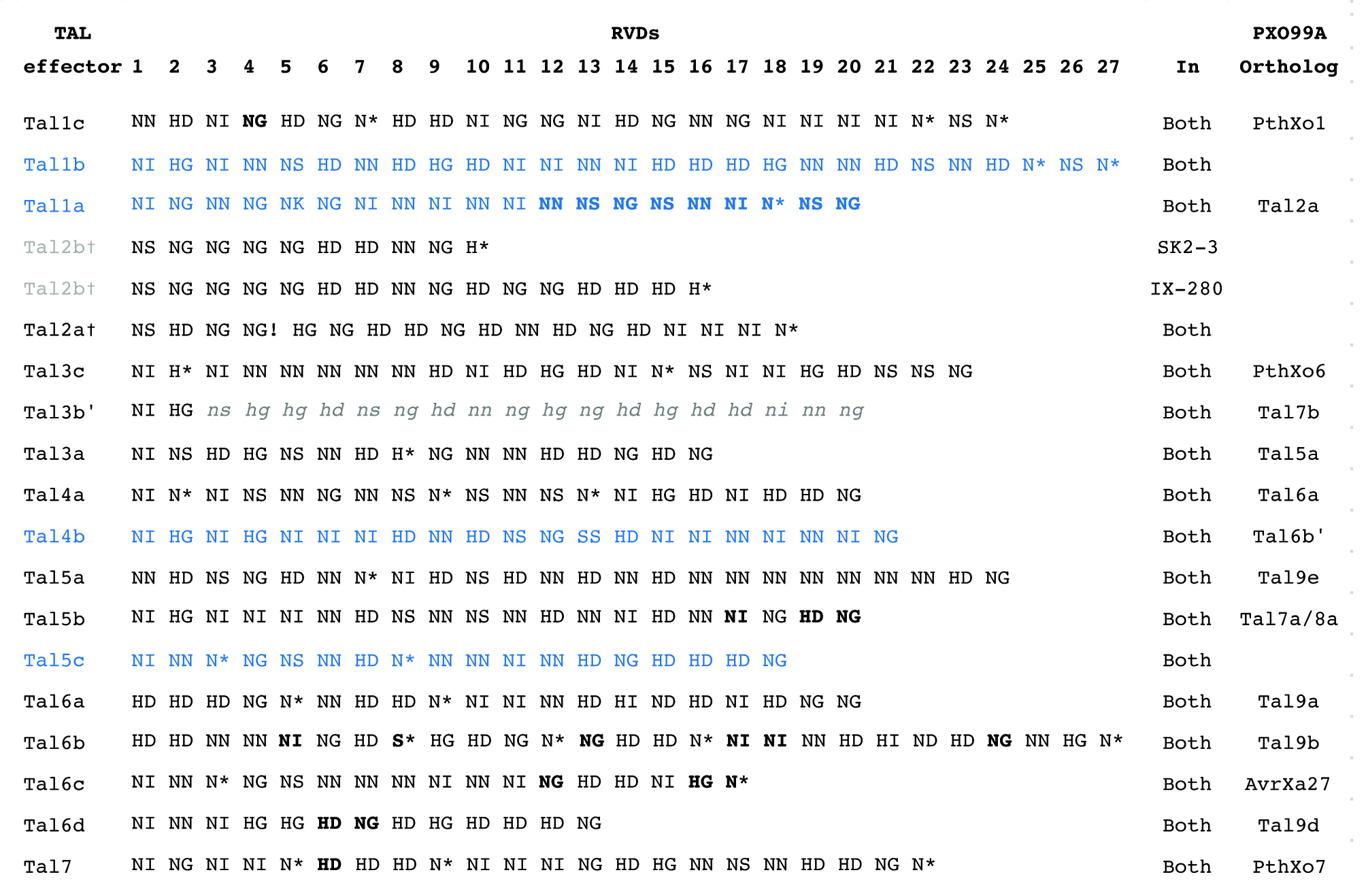
RVD sequences of IX-280 and SK2-3 TALEs. RVDs in bold are different in PXO99A orthologs. A dagger indicates a truncTALE. Grey font indicates TALEs that are not the same in IX-280 and SK2-3. Lower case italicized RVDs are untranslated following a frameshift. Blue font highlights TALEs for which EBEs in rice were predicted.

Tal1c of both strains is an ortholog of PthXo1 (Figure 4), which likely explains the ability of each strain to overcome *xa5.* PthXo1 in IX-280 and SK2-3 differs from PthXo1 in PXO99A at one RVD, but the base-specifying residue of that RVD is the same (Supplementary Figure S3). Notably, an ortholog of PthXo7, the PXO99A TALE that induces *TFIIAγ1*, is also present in both strains (Tal7). Compatibility with *xa5* had been postulated to be due to activation of the paralog *TFIIAγ1* by PthXo7 ^45^, but it was recently shown that only TFIIAγ5, and not TFIIAγ1, interacts *in planta* with tested TALEs ^46^.

TALEs that could enable defeat of *xa13* are less apparent. IX-280 and SK2-3 have no ortholog of known, major virulence factors such as PthXo2, which drives expression of *SWEET13* (also called *Os12N3* or *Xa25*), or of PthXo3, TalC, Tal5, or AvrXa7, which activate *SWEET14* ^7,8,47-49^. In all, eleven of the 15 TALEs of IX-280 and SK2-3 (including Tal1c and Tal7) are apparent orthologs of TALEs found in PXO99A, which does not overcome *xa13* (Figure 4, Supplementary Figure S3). Of these, six are identical to their PXO99A counterpart and the others have from one to several differences in RVD sequence. The gene encoding Tal4b of IX-280 and SK2-3 has an ortholog in PXO99A that is pseudogenized by a frameshift early in the coding sequence. We reason that the *xa13*-compatibility of IX-280 and SK2-3 is conferred by one or more of the TALEs with no apparent, intact ortholog in PXO99A (i.e., Tal1b, Tal1a, Tal4b, and Tal5c] or with a difference in RVD sequence relative to the PXO99A counterpart (i.e., Tal1c, Tal5b, Tal6b, Tal6c, Tal6d, and Tal7).

### 2.5. Predicted targets of possible *xa13*-breaking TALEs

Toward identifying the basis for IX-280 and SK2-3 compatibility with *xa13*, we generated lists of candidate target genes in rice (cv. Nipponbare) for their Tal1a, Tal1b, Tal4b, and Tal5c, which are either not found in PXO99A or differ by more than 6 RVDs from the most similar TALE in PXO99A (see Materials and Methods). Tal1a contains several instances of RVDs NN and NS, which have dual and lax specificity, respectively, so EBEs were predicted in most promoters. Among the candidates for Tal1b was a *SWEET* gene, *SWEET2b*, but it was shown previously not to function as an *S* gene ^7^. Another was a putative sulfate transporter gene. The putative sulfate transporter gene *OsSULTR3;6* is an *S* gene for bacterial leaf streak caused by *X. oryzae* pv. oryzicola ^50^, but whether sulfate transporters might confer susceptibility in bacterial blight is unknown; heterologous expression of an *OsSULTR3;6*-inducing TALE in the TALE-deficient strain X11-5A did not increase the extent of bacterial blight caused by this strain ^51^. For Tal4b, EBEs were predicted in the promoters of three *SWEET* genes, *SWEET1b* (clade I and shown not to function as an S gene ^7^), *SWEET7e* (clade II and not tested), and *SWEET14*, within 350 bp of the transcriptional start sites (TSS). For Tal5c, EBEs were predicted in the promoters of *SWEET15* within 100 bp of the TSS, *SWEET13* within 250 bp, and *SWEET12* within 50 bp.

### 2.6. IX-280 compatibility with *xa5* is associated with induction of *SWEET11* but IX-280 induces no clade III *SWEET* in compatible *xa13* plants

Testing candidate targets of each of the IX-280 and SK2-3 TALEs that differ from TALEs of PXO99A to determine the mechanism by which these strains overcome *xa13* was beyond the scope of this study. We focused instead on just the clade III *SWEET* genes, including *SWEET12*, which was not a predicted target. We inoculated IX-280 to rice cultivar IR24, which harbours neither *xa5* nor *xa13*, and near isogenic cultivars IRBB5 (*xa5*), IRBB13 (*xa13*), and IRBB53 (*xa5* and *xa13*). Each of these cultivars except IRBB53 is susceptible to IX-280 (^21^ and Figure 5a). We hypothesized that IX-280, by virtue of its PthXo1 ortholog Tal1c, induces *SWEET11* strongly in IR24 and sufficiently in IRBB5, and that for compatibility in IRBB13 it induces another *SWEET* gene or the *xa13* allele of *SWEET11* by virtue of some other TALE. Further, we hypothesized that induction of the alternate *SWEET* gene is not as strong as that of *SWEET11*, such that when diminished by *xa5* it is insufficient for susceptibility, explaining incompatibility with the combined *xa5* and *xa13* rice genotype IRBB53. We first compared expression of *SWEET11* across each of the cultivars, using quantitative RT-PCR of RNA harvested from leaf tissue 24 hr after inoculation. It was induced to 799 fold in IR24, to 553 fold in IRBB5, and not at all in IRBB13 or IRBB53, relative to mock (water) inoculation (Figure 5b). For reference we also examined expression of the bZIP transcription factor gene *TFX1* and the *TFIIAγ*5 paralog *TFIIAγ*1. These are targets of PXO99A TALEs PthXo6 and PthXo7; these TALEs contribute moderately to virulence ^45^ and an ortholog of each (Tal3c and Tal7, respectively) is present in IX-280 and SK2-3. In IR24 and IRBB13 each of the transcription factor genes was moderately induced (20-35 fold) in IX-280-inoculated leaves relative to mock (Figure 5b). This induction provides evidence that Tal3c and Tal7 are delivered and functional, and that the single RVD difference between PthXo7 and Tal7 does not impact targeting of *TFIIAγ1*. In IRBB5 and IRBB53, *TFX1* and *TFIIAγ*1 induction was reduced to just 3-5 fold relative to mock (Figure 5b). This result is consistent with the observation that the *xa5* allele reduces generally the ability of TALEs to induce their targets ^46^. Next, we assayed the ability of IX-280 inoculated to IRBB13 plants to induce any of the other clade III *SWEET* genes. Contrary to our hypothesis, it induced none (Figure 5c).

**Figure 5:**
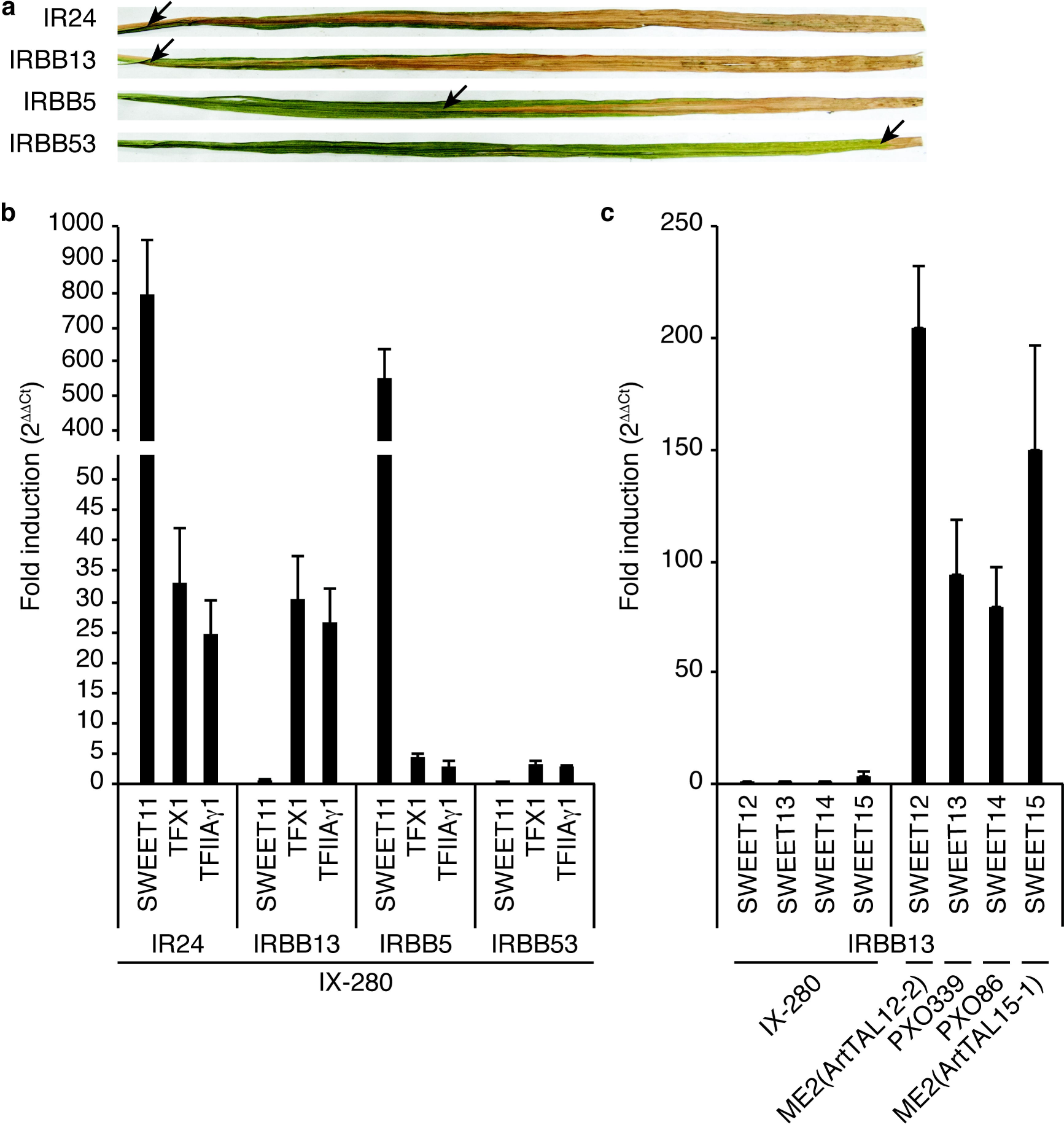
Compatibility and ability of IX-280 to induce known or potential bacterial blight *S* genes in near-isogenic rice lines IR24, IRBB5 (*xa5*), IRBB13 (*xa13*), and IRBB53 (*xa5* and *xa13*). (a) Representative lesions at 14 days after clip inoculation. Arrows indicate the distance the lesion progressed. (b) Fold induction of *SWEET11*, *TFX1*, and *TFIIAγ1* 24-27 hr after inoculation by syringe infiltration of IX-280 relative to mock (water)-inoculated leaves, measured by qRT-PCR. Each bar represents the mean of three replicates. Error bars represent standard deviation. (c) Fold induction, as in (b), of the other clade III *SWEET* genes by IX-280 and selected positive control strains. ME2 is a *pthXo1* knockout derivative of PXO99A ^6^ used here to deliver artificial TALEs ArtTAL12-2 and ArtTAL15-1, which are targeted to the *SWEET12* and *SWEET15* promoters, respectively ^7^. PXO339 is a Philippines race 9 *Xoo* strain that induces *SWEET13* ^48^. PXO86 is a Philippines race 1 *Xoo* strain that induces *SWEET14* ^8,55^.

## 3. Discussion

This study presents the first completely assembled genome sequence of an Indian *Xoo* strain and the first genome sequence of a Thai strain. The genome comparisons we carried out (Figure 2a) and comparisons published elsewhere ^42,52^ demonstrate the high level of variability in genome structure across different strains of *Xoo* and a general lack of relationship between genome structure and the geographical location at which a strain was isolated. Like other *Xoo* strains, both IX-280 and SK2-3 contain hundreds of IS elements and other transposons in their genomes (Table 1) that likely contribute to genome plasticity ^24,52^. Despite the overall genome structure variability in the species and the geographic separation of IX-280 and SK2-3, strikingly these two strains are part of a young and highly clonal lineage prevalent in India, L-I ^19^, that contains no other characterized, non-Indian strains. This observation and the relative rarity of *xa*5 compatibility in Thailand (Figure 1b) suggest introduction of SK2-3 or a recent progenitor in lineage L-I to Thailand from India. Alternatively, members of the lineage could have been introduced separately to India and to Thailand.

**Table 1:** The IX-280 and SK2-3 genome assemblies.

|  | **IX-280** | **SK2-3** |
| --- | --- | --- |
| **Chromosome** | **4,963,593 bp** | **4,934,446 bp** |
| **Plasmid** | **42,975 bp** | **-** |
| **Final coverage** | **164.0x** | **156.4x** |
| **% Mapped reads** | **94.6%** | **92.0%** |
| **Annotated genes** | **5,041** | **4,926** |
| **Annotated IS elements** | **411** | **407** |
| **Annotated transposases** | **730** | **698** |
| ***tal* genes** | **17** | **17** |

The strains IX-280 and SK2-3 are of special interest because of their compatibility with multiple single *R* genes ^21,22^, in particular *xa5* and *xa13*. Previously, strains compatible with *xa5* and with *xa13* were only found in the genomically and geographically diverse lineage L-3 ^19^; IX-280 and SK2-3 represent the first example of strains compatible with *xa5* and with *xa13* in the much more genetically homogeneous lineage L-I. In light of the expanding *R* gene compatibility and geographic spread of strains in L-I, the fact that IX-280 and SK2-3 are incompatible with the *xa5* and *xa13* stacked line IRBB53 underscores the potential benefit of deploying such *R* gene stacks. The results also illustrate the importance of complete genome sequencing in monitoring *Xoo* populations to develop and deploy varieties with effective disease resistance.

The basis for the compatibility of IX-280 and SK2-3 with *xa5* is almost certainly their ability to sufficiently activate *SWEET11* even under the dampening effect of *xa5* (Figure 5b). Their PthXo1 ortholog, Tal1c, is presumably responsible for this; the single difference in RVD sequence between Tal1c and PthXo1 does not affect the base specifying residue (Supplementary Figure S3). Induction of *TFX1* by Tal3c (the PthXo6 ortholog), and of *TFIIAγ1* by Tal7 (ortholog of PthXo7), though reduced by *xa5*, may also contribute. As noted, PthXo6 is a demonstrated virulence factor and *TFX1* is a verified *S* gene ^45^. PthXo7 is also a demonstrated virulence factor, and although activation of *TFIIAγ1* was observed only by the *xa5*-compatible strain PXO99A ^45^, silencing it decreased susceptibility to PXO99A even in an *xa5* background ^46^. We also observed that despite induction of *TFIIAγ*1 by Tal7, activation of *SWEET11*, *TFX1*, and *TFIIAγ*1 itself remain dampened in IRBB5 relative to IR24 and IRBB13 (Figure 5b). Thus, activation of *TFIIAγ1* by Tal7 appears to contribute to susceptibility in some way other than providing a substitute for *TFIIAγ5*.

The basis for the compatibility of IX-280 and SK2-3 with *xa13* is yet to be determined. Despite some of their TALEs being predicted to target clade III *SWEET* genes, no clade III *SWEET* gene was induced by IX-280 in IRBB13 plants (possible reasons for such false positive predictions include competing endogenous DNA-binding proteins or DNA methylation at the target, or binding that does not lead to gene activation due to position in the promoter). Compatibility with *xa13*, which harbours a promoter deletion that eliminates the binding site of PthXo1 (and of the IX-280 and SK2-3 ortholog Tal1c), but not with *xa5* and *xa13* together, thus points to a second major TALE in IX-280 and SK2-3 that activates an alternative, novel *S* gene. Because of the incompatibility with stacked *xa5* and *xa13*, such as in IRBB53, one would predict that the induction of this alternative *S* gene by the TALE is not strong enough to remain effective when dampened by *xa5*. Though we predicted targets only for IX-280 (and SK2-3) TALEs most dissimilar to those of the *xa13*-incompatible strain PXO99A, it is possible that one of the IX-280 TALEs more closely related to a PXO99A TALE is responsible: even a single RVD difference could confer the ability to target a new gene. It is also possible that the promoter sequence of the alternative *S* gene is different in IRBB13 and not represented in our predictions using the Nipponbare reference. Thus, future work should begin with loss- and gain-of-function experiments for each of the IX-280 and SK2-3 TALEs that are not precisely conserved in PXO99A, followed by transcript profiling of the IRBB13 and IR24 responses for any unique TALE revealed to be important for compatibility in IRBB13 plants.

Studies of diverse strains have suggested that induction of a clade III *SWEET* gene is a fundamental requirement for *Xoo* to cause bacterial blight of rice, but the African *Xoo* strain BAI3 was recently reported to be compatible on a rice line from which the binding site for its major TALE, TalC, in the promoter of *SWEET14*, was removed by genome editing, and the strain induced no clade III *SWEET* in that line ^53^. We have shown for the first time compatibility of an Asian *Xoo* strain without clade III *SWEET* gene induction. The emerging picture suggests some degree of selection on *Xoo* populations to evolve to target alternative *S* genes, perhaps due to the extensive deployment of *R* genes like *xa13* and *xa25* (a recessive allele of the *SWEET13*/*Xa25 S* gene widely used in China) ^12,48,54^. It will be of interest to determine whether the evolutionarily divergent IX-280 (and SK2-3) and BAI3 lineages have converged on the same new *S* gene, or if they target distinct ones. Further, determining the biochemical function(s) of the new *S* gene(s), which ostensibly can substitute for sucrose export in rendering the plant susceptible, promises to shed light on the mechanism by which clade III *SWEET* genes contribute to disease development.

## Acknowledgements

This work was supported by the Indian Council of Agricultural Research-Networking Project on Transgenic Crops (ICAR-NPTC) and Department of Biotechnology (DBT) (award BT/CEIB/12/1/01 to RR) and by the U.S. National Science Foundation (award 1444511 to AJB.) Support from the National Phytotron Facility (NPF) at the Indian Agricultural Research Institute (IARI), New Delhi, India is gratefully acknowledged.

## Authors’ contributions

GSL, RO, SP, and RR conceived the study; SCDC, AJB, and RR designed the experiments; SCDC, PM, CG, PD, LW, SM, WK, JSL, and RR performed the experiments and/or generated data; SCDC, LW, PBP, NKS, KS, AJB, and RR analysed and interpreted the data; SCDC, AJB, and RR drafted the manuscript with input from RO.

## Conflict of interest

The authors declare no conflict of interest.

